# The transcription regulatory code of a plant leaf

**DOI:** 10.1101/2020.01.07.898056

**Authors:** Xiaoyu Tu, María Katherine Mejía-Guerra, Jose A Valdes Franco, David Tzeng, Po-Yu Chu, Xiuru Dai, Pinghua Li, Edward S Buckler, Silin Zhong

## Abstract

The transcription regulatory network underlying essential and complex functionalities inside a eukaryotic cell is defined by the combinatorial actions of transcription factors (TFs). However, TF binding studies in plants are too few in number to produce a general picture of this complex regulatory netowrk. Here, we used ChIP-seq to determine the binding profiles of 104 TF expressed in the maize leaf. With this large dataset, we could reconstruct a transcription regulatory network that covers over 77% of the expressed genes, and reveal its scale-free topology and functional modularity like a real-world network. We found that TF binding occurs in clusters covering ∼2% of the genome, and shows enrichment for sequence variations associated with eQTLs and GWAS hits of complex agronomic traits. Machine-learning analyses were used to identify TF sequence preferences, and showed that co-binding is key for TF specificity. The trained models were used to predict and compare the regulatory networks in other species and showed that the core network is evolutionarily conserved. This study provided an extensive description of the architecture, organizing principle and evolution of the transcription regulatory network inside the plant leaf.

## Introduction

For all living cells, a basic control task is to determine gene expression throughout time and space during developmental processes and responses to stimuli^1^. The code that governs gene expression is stored in the non-coding region of the genome. Sequence-specific DNA binding proteins known as transcription factors (TFs) can read the code by binding to cis-regulatory elements to activate or repress gene expression. Each gene can receive instructions from multiple TFs, and each TF can target thousands of genes, forming a transcription regulatory network, which regulates almost every biological processes inside the cell^2^. Hence, it is critical to understand the mechanism, architecture and behaviour, as well as conservation and diversification of the regulatory network.

The budding yeast has probably the best mapped transcription regulatory network and is a unicellular model for eukaryote, with around 200 TFs^3,4^. For multicellular organisms with large genomes and thousands of TFs, complete network reconstruction has proven to be a formidable task. For example, the ENCODE consortium has mapped the binding of 88 TFs in five cell lines to study its architecture and dynamics^5^, while a more complete regulatory network was constructed in human colorectal cancer cells with data from 112 TFs^6^. But similar efforts are seldom feasible for individual laboratories, and have yet to be attempted in plants.

Maize is one of the most economically important crops worldwide. It is also the best-studied and most tractable genetic system among the cereal crops, making it an ideal model for studying this group of important crop plants^7^. It has been shown that the maize non-coding region harboring the cis-regulatory elements is a major contributor to the observable phenotypic differences and adaptation, between and within species. For example, 70% of maize GWAS hits are located in the non-coding regions that has yet been functionally annotated^8^. Recently, it was shown that open chromatin sequence variations within these non-coding regions could explain up to 40% of the phenotypic variations of key agronomic traits^9^. However, it is difficult to assess their true functions without knowing the precise locations of the cis-regulatory elements and the TFs that recognize them. Despite the importance of TF binding data, ChIP-seq experiments in plants are limited by antibody availability and difficulties in transforming crops to express the epitope fusion protein. As a result, TF binding information is only available for a handful of maize TFs, and even in the model species like Arabidopsis and rice, such studies are too few in number to produce a general picture of the plant transcription regulatory network.

In this study, we used 104 TF ChIP-seq to reconstruct the maize leaf transcription regulatory network, and trained machine-learning models to predict TF binding and co-localization, and annotate non-coding regions in other grasses. Our findings provide a first glimpse into the plant transcription regulatory network and reveals its architecture, organizing principle and evolutionary trajectory.

## Results

### High-throughput ChIP-seq using maize leaf protoplast

We have developed an efficient protoplast isolation and transformation method that enabled us to expressed TF constructs for high-throughput ChIP-seq (Fig. 1a, Methods). With it, we have successfully performed 218 ChIP-seq experiments for 104 TFs that are expressed in the developing section of the maize leaf based on previous RNA-seq data^10^ (Fig. 1a and Supplementary Table 1). Next, we used the ENCODE2 uniform pipeline to process the ChIP-seq data^11^ (Fig. 1a and Supplementary Fig. 1a). We identified a total of 2,147,346 reproducible TF binding peaks, and the number of peaks varies between TFs, with a median value of ∼16K (Interquartile range, between 7,664-32,566 peaks; Supplementary Fig. 1b). We anticipated that plant TF binding might form dense clusters and frequently locate within open chromatin regions based on previous mammalian studies^6^. To test this, we merged all TF binding peaks in the genome and found that they indeed clustered into 144,890 non-overlapping loci, covering ∼2% of the genome (Supplementary Table 2). Similar saturation of TF binding clusters was observed when sufficiently large number of TFs expressed in colorectal cancer cells have been sampled^6^.

**Fig. 1.**
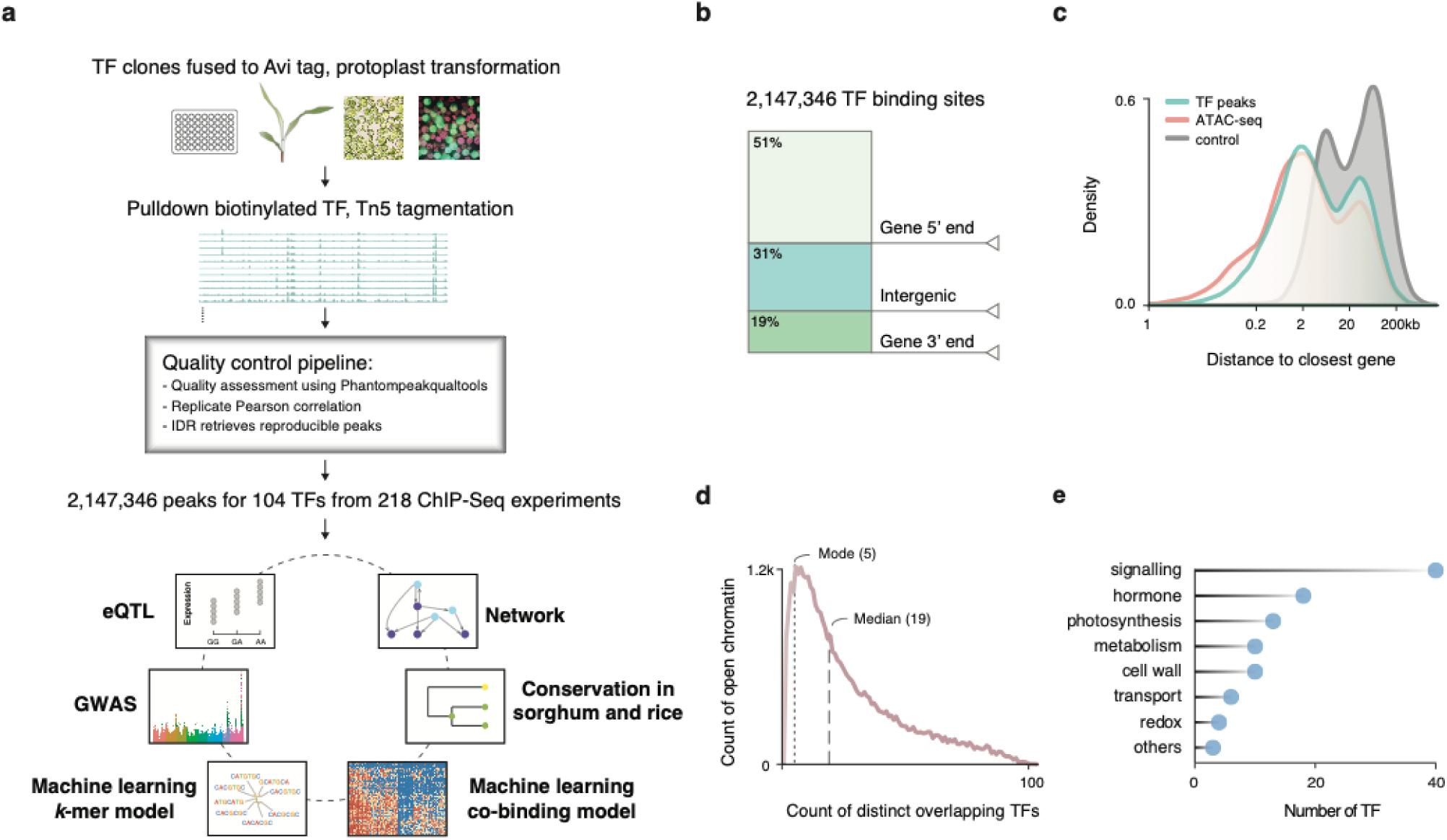
The regulatory landscape of the maize leaf. **a**, Overview of the experimental and analytical approaches. **b**, Genome-wide distribution of TF binding loci. **c**, Density plots corresponding to distances of TF binding sites (green) and open chromatin regions (red) to closest annotated gene. Gene to gene distant is used as a control (grey). **d**, Distribution of the count of distinct TFs that overlap with open chromatin regions. **e**, Functional classification of the 104 TF based on their target genes.

Next, we measured chromatin accessibility in the same tissue using ATAC-seq, and identified 38,713 biologically reproducible open chromatin regions. We found that the TF binding loci and open chromatin regions show similar genome-wide distributions, frequently proximal to gene bodies (+/- 2.5 kb), with preferences for the 5’ end (Fig. 1b,c). We also found that the distance between TF binding site to gene after excluding regions that overlap with gene body is bimodal (Fig. 1c). Despite the larger space available for distal regulation, we observed that regions between 10-100 kb constitute ∼15% of the TF binding loci, and ∼17% of the open chromatin regions.

Layering ATAC-seq and ChIP-seq data showed that TF binding loci and open chromatin overlap (Supplementary Fig. 1c; P-value < 10^−5^). On average, ∼74% of the peaks for a given TF (IQR_25-75_ 64%-87%) intersect with open chromatin regions (Supplementary Fig. 1d) confirming the relevance of the identified TF binding sites within the chromatin context. Collectively, 98% of the open chromatin regions overlap with TF peaks (Fig. 1d), with a mode of 5, and a median of 19 distinct TFs for each region, suggesting there are a large number of possible TF combinations that co-regulate transcription, in contrast to the classic view of a singular or few regulators that control the expression of genes.

As none of the 104 TFs selected have been previously examined by ChIP-seq, and most of them have not even been functionally characterized, we used GO-term and MAPMAN functional category enrichment analysis to classify them based on their targets (Fig. 1e and Supplementary Table 3). Majorities of the TFs are grouped into signaling, hormone, photosynthesis and metabolism categories, which are the core biological functions of the leaf. For example, targets of the maize ZIM TFs show enrichment in the GO terms “response to wounding” and “jasmonic acid metabolism”, consistent with the role of their homologs in other plant species^12^. The PIF and GLK TFs targets are enriched for “circadian rhythm”, “light harvesting” and “apocarotenoid metabolic process”, and were assigned to the photosynthesis category. The *ZmMyb38*/*ZmCOMT1* targets were associated with terms such as “regulation of flavonoid biosynthetic process”, and “phenylpropanoid biosynthetic process” and were assigned to the metabolic group^13^.

Previous studies showed that the high TF occupancy regions in the genome are often associated with important functions^14,15^. We identified 2,037 open chromatin regions at the top 5% of the TF occupancy distribution (Supplementary Table 4), and their surrounding genes are indeed enriched for regulatory GO-terms (Supplementary Table 5). Notably, we found that *Vgt1*, the most important QTL for flowering time^16^, is associated with a high TF occupancy region bound by 76 TFs, and locates at a distance of ∼72 kb from the *ZmRAP2*.*7*, a TF gene whose expression is regulated by *Vgt1*. Six of the TFs bound to this region (*i*.*e*., PRR5, ELF3, COL3, COL7, COL18 and DOF3/PBF1) have been linked to flowering time variations through previous genetic studies^17,18^.

Taken together, our data provide a comprehensive catalogue of the biological functions of maize TFs. Interestingly, despite over 1 billion years of divergence, TF binding formed dense clusters in plant and animal genomes, and high occupancy regions play an important role in shaping complex phenotypic variations.

### TF binding sites show sequence conservation and enrichment for eQTL and GWAS hits

TF binding is a key determinant of transcriptional regulation and, if purifying selection is effective in these regions, they should exhibit low sequence diversity. We examined the conservation of the TF binding sites by assessing the overall nucleotide diversity represented in the maize HapMap^19^ while controlling for overall SNP density in function to the distance of TF’s peak summit (Fig. 2a). The result confirmed that sequence variation is, in fact, reduced.

**Fig. 2.**
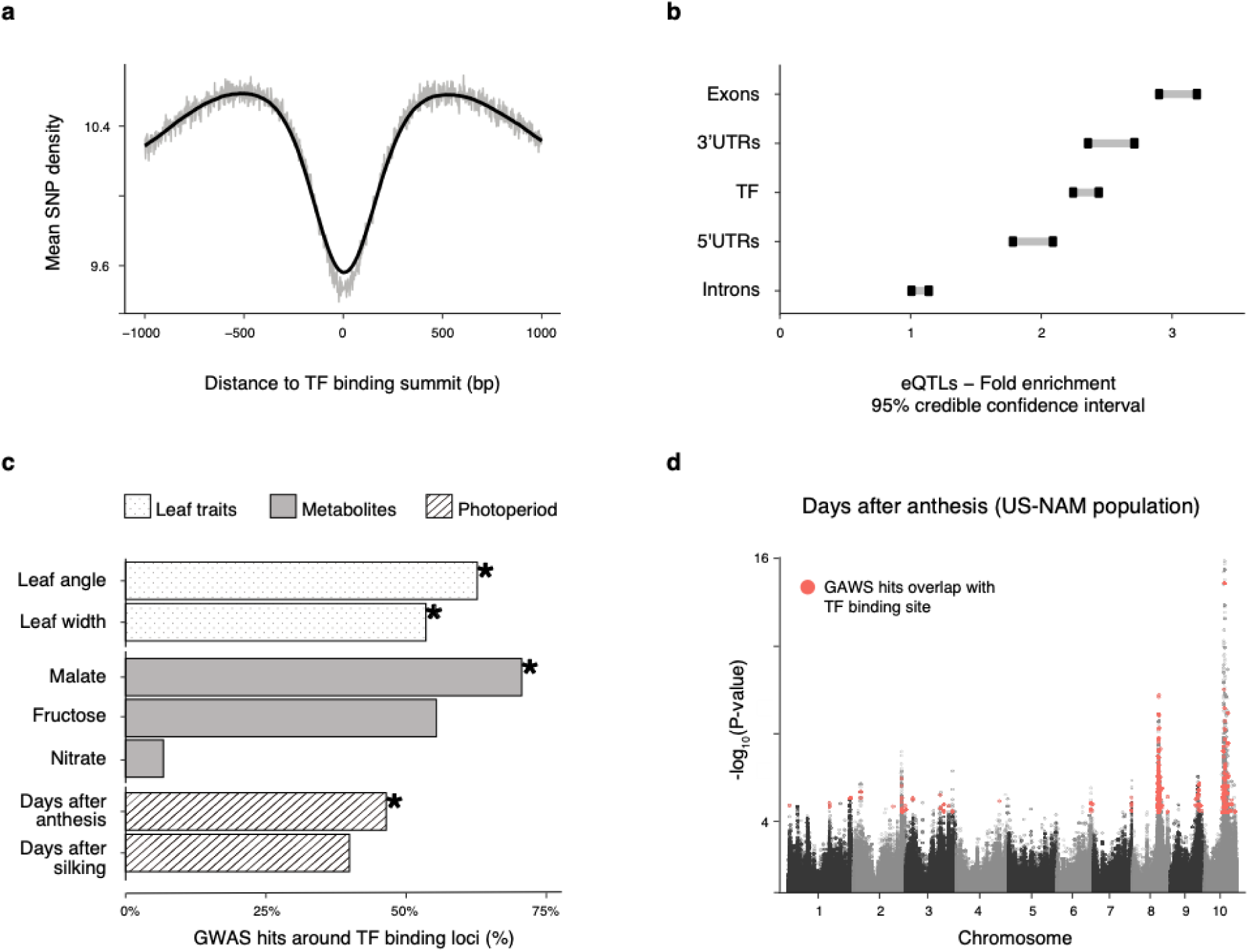
TF binding sites show low sequence diversity and enrichment in functional variation. **a**, Mean SNPs density calculated in sliding windows of 100 bp bins flanking TF binding summits, and normalized by mappability rate. **b**, 95% confidence interval for the enrichment of SNPs associated to variation in mRNA levels (eQTLs) vs. non eQTLs SNPs, and relative to control regions for different sets of genomic regions. **c**, Proportion of phenotype-associated GWAS hits for an assortment of traits overlapping to TF binding loci. Traits in which the enrichment was statistically significant are labelled with an asterisk. **d**, Manhattan plot of GWAS for days after anthesis. Highlighted GWAS hits overlap with binding regions for a group of TFs (PRR5, ELF3, COL3, COL7, COL18 and DOF3/PBF1) associated to photoperiod variation.

While binding of most TFs is constrained, TF and their binding sites are key to local adaptation or domestication. For example, TF *ZmTB1* is known to play an important role in maize domestication^20^. Hence, we predict that TF binding loci could be enriched for common SNP variations controlling gene expression and downstream traits. This was first tested in a panel of 282 inbred breeding lines for their effect on mRNA expression using common and likely adaptive variants^21^. We found two-fold enrichment of TF binding loci (95% credible interval 2.26-2.46), similar to the enrichment around 5’ and 3’ UTRs (Fig. 2b). Distal and proximal TF binding loci are also enriched when examined separately (Supplementary Fig. 2a). The enrichment pattern is ubiquitous across the 104 TFs that we studied (Supplementary Fig. 2b and Supplementary Table 6). Overall, these support our hypothesis that common variation in mRNA expression is controlled by TF binding site variation.

Next, we tested their overlap with functional variations associated with agricultural traits others than gene expression. To this end, we calculated the enrichment in GWAS hits for seven traits related to metabolites^22^, leaf architecture^23^, and photoperiodicity^24^ measured in the US NAM population. Overall, TF binding loci are enriched for four of the traits (Fig. 2c, Supplementary Table 7). We noticed that simple traits, such as metabolites, showed few TFs enriched for GWAS hits (*e*.*g*. malate and nitrate). In contrast, many TF bindings overlap with GWAS hits for complex traits, which are known to be polygenic and influenced by a large number of genetic variants (*e*.*g*., days after silking and days after anthesis; Fig. 2d and Supplementary Fig. 2c). A further look at TFs enriched in GWAS hits for photoperiodicity revealed that 51% of the TFs enriched in days after anthesis, and 35% in days after silking, also bound to the *Vgt1/ZmRAP2*.*7* high TF occupancy region, suggesting they could be novel regulators of maize flowering time.

Overall, our observations that TF binding regions are conserved and frequently overlap with sequence variations associated with phenotypic changes meet our initial expectations. The general trend for GWAS enrichment also supports our hypothesis that non-coding region variations related to traits are mediated by TFs. Furthermore, our finding highlighted the potential of using large-scale TF ChIP-seq data to connect sequence variation in *cis* to *trans*-regulators to highlight the molecular mechanism implicated in complex phenotypes.

### A scale-free transcription regulatory network in plant

Real-world networks such as the www, social network, human gene regulatory network and protein-protein interaction network, frequently exhibit a scale-free topology^25^. To assess the feasibility of our data to pinpoint true regulatory relationships, we tested if the derived network would fit this topology. We reshaped the regulatory data into a graph (Fig. 3a), adopting the ENCODE probabilistic framework to identify high confidence proximal interactions (P-value < 0.05)^5,11,26^. This approach renders a network with 20,179 nodes (∼45% of the annotated genes and ∼77% of the leaf expressed genes, Supplementary Table 8). We evaluated the in-degree distribution (Fig. 3b), or number of edges towards each node, and found it to follow a linear trend in the log-scale (R^2^ = 0.882, P-value < 2.2e-16), as expected for power-law distribution (goodness of fit P-value = 0.67), which is a landmark of scale-free networks.

**Fig. 3.**
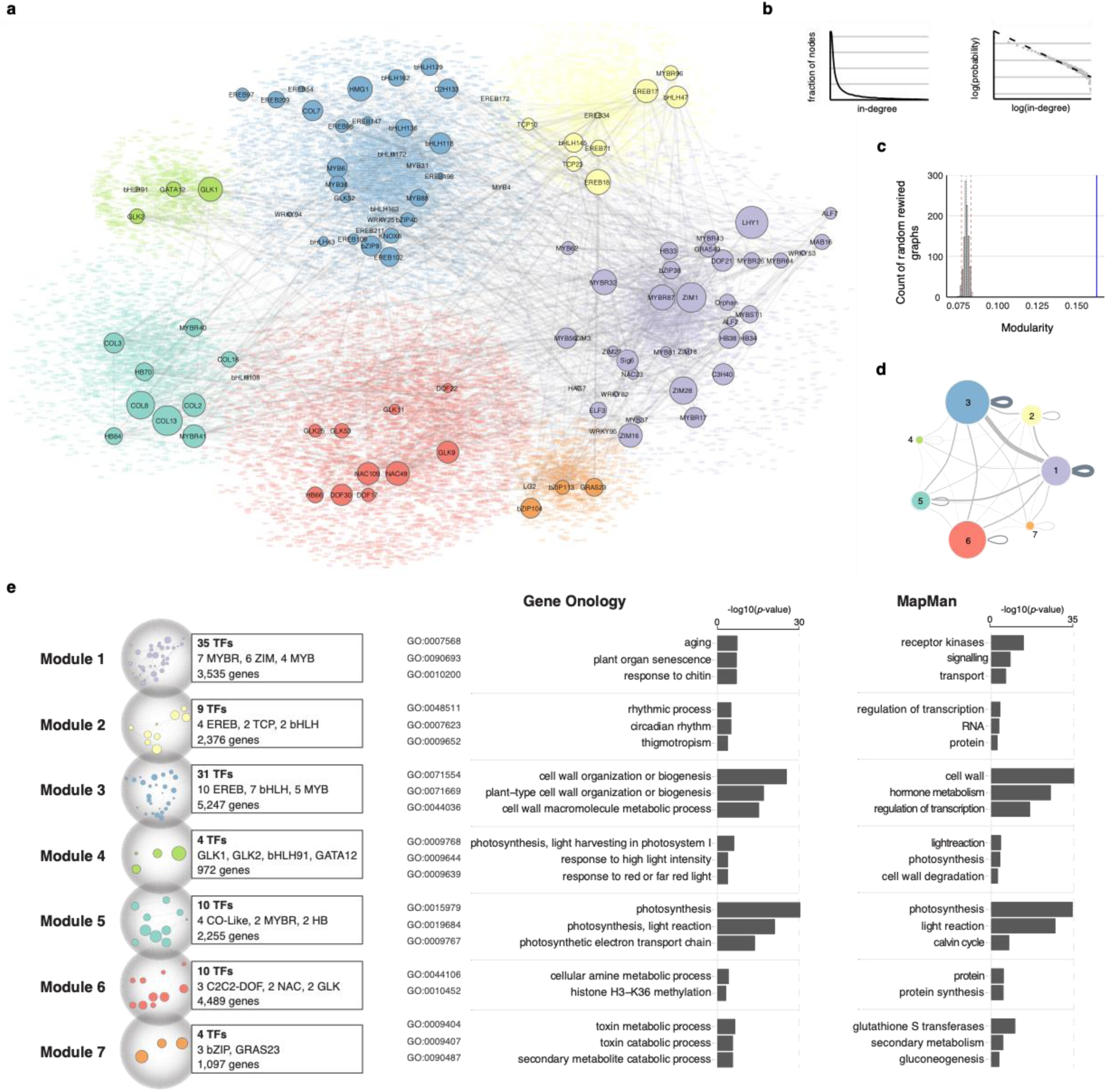
The maize leaf transcription regulatory network exhibit properties of real-world networks. **a**, Graph diagram showing the maize transcription regulatory network. The seven modules are labelled in different colors. TF nodes are shown as circles with size representing gene expression level, and their target genes are shown as color dots in the background. **b**, Distribution of node in-degree values shows it fits the power-law (dashed line). **c**, Modularity of the network (blue line) is higher than the random control (histogram of maximum modularity values of 1000 rewired graphs). **d**, Graph diagram showing TF to gene connections within and between modules. Each module is represented as circle with size proportional to the number of nodes within. **e**, GO-term and MapMan category enrichment analysis of genes in each module.

In a real-world network, nodes that appear more connected than others are called “hubs”, and they are critical for information flow^25^. We defined these hub genes as target nodes in the top percentile of the in-degree distribution (99^th^ percentile, for a total of 206 genes). Interestingly, we found half of the hub genes were located nearby the TF highly occupied regions (Supplementary Table 5). Similar to the result from the analysis of highly occupied regions, hub genes showed enrichment for regulatory functions (*i*.*e*., GO term: “*transcription factor activity*”, enrichment 4-fold), consistent to the expected role for a hub node in the transcription regulatory network.

### Structural and functional modularity of the network

Biological networks often exhibit topological and/or functional modularity^27,28^. We tested for this predicted property by contrasting the maximum modularity in the network to a null distribution from an ensemble of random rewired graphs (H_0_: 1000 rewired graphs)^29^. The result confirmed that the network exhibits a significant increase in modularity (P-value < 0.05, Fig. 3c). Next, we applied the partitioning algorithm in Gephi to determine relationships between subsets of network elements at different scales. At resolution 1.8, the whole graph will be collapsed into a single module, and at resolution of 0 every node will be one module. Between the two extremes, at resolution 1, the network can be divided into seven modules, each ranging from ∼27% to ∼5% of the total nodes (Supplementary Table 9). These modules are not isolated. Only ∼40% of the total edges occurring within each module, suggesting that TFs often regulate targets outside their own modules, and there are large information flows between modules (Fig. 3d).

We predicted that topological modularity could be related to function in known biological processes. Hence, we performed GO-term and MapMan functional category enrichment analyses for genes in each module, and found that they were indeed enriched for specific functions (Fig. 3e). For example, we found two photosynthesis-related modules. Module 4 is enriched in targets of GLK1/GLK2, which are known regulators of photosynthesis^30^, while model 5 is enriched in targets of the CONSTANT-like (COL) TFs, which are associated with roles in flowering time, circadian clock and light signaling^31^ (Supplementary Table 10).

These modules contain thousands of genes from different pathways, and are too large to be assessed as a whole. We hypothesize that as the network topology can already provide clues to biological function at this scale, potential regulators of a smaller pathway might be identified based on connectivity at a local scale. We first tested this in the conserved chlorophyll biosynthesis pathway, which is known to be regulated by two GLK TFs, as their mutations caused disruption of the photosynthesis apparatus gene expression^30,32^. We calculated the sum of the log transformed p-values that the ENCODE probabilistic model generated for each TF-target interaction based on binding intensity and proximity, and used it to infer the contribution of each TF to the pathway (Fig. 4a). We found that the top regulators are indeed the two GLKs and an unknown MYBR26. Despite the function of the MYBR26 has yet been studied, its Arabidopsis homologs are involved in circadian regulation, confirming our hypothesis^33^.

**Fig. 4.**
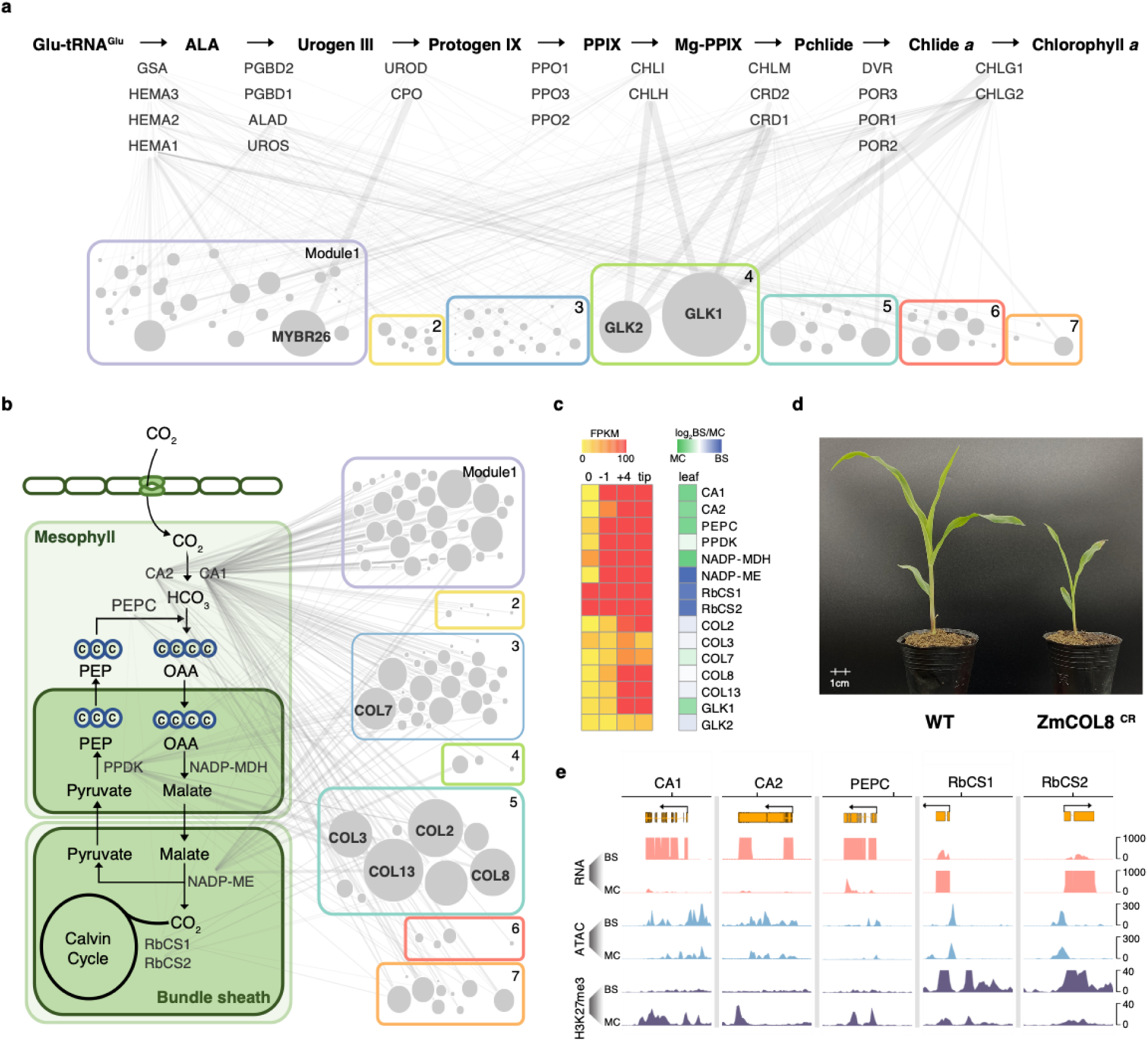
Identify key regulators based on local TF connectivity. **a**, The chlorophyll biosynthesis pathway. The genes encoding key enzymes for each step are shown below the arrow. The width of the edge between TF and gene represent regulatory potential based on the TIP model. The 104 TF nodes are represented as circles and grouped into 7 modules, with size proportional to the sum of the regulatory potential. **b**, The maize photosynthesis pathway. Key genes encoding enzymes responsible for CO_2_ fixation and transport between mesophyll and bundle sheath cells are shown. **c**, Heatmap shows the expression gradient of core C_4_ photosynthesis genes in four leaf sections from base to tip, as well as their mesophyll and bundle sheath cell-specific expression pattern. **d**, Pale-green leaf and seedling lethal phenotypes of the COL8 CRISPR/Cas9 mutant. **e**, Cell-type specific H3K27me3 in core C_4_ gene loci. MC: mesophyll cell; BS: bundle sheath cells.

Next, we used this strategy to examine the maize C_4_ photosynthesis pathway without well-defined regulator. It turns out that the top five TFs in connectivity ranking are COL TFs (Fig. 4b,c). Previous studies of COLs in other plant species have showed that they play an important role in the regulation of flowering and photoperiod^31^. Without maize COL mutants, we searched different maize CRISPR/Cas9 populations and found one line with a frame-shift deletion in the first exon of *COL8* (Supplementary Fig. 3). The homozygous mutant has a pale-green and seedling lethality phenotype, supporting our hypothesis that the COL TFs are important for photosynthesis (Fig. 4d). Interestingly, for key C_4_ photosynthesis genes that are expressed specifically in mesophyll or bundle sheath cells, we found that their loci are associated with cell-specific H3K27me3 mark, suggesting that they are regulated not just by a complex TF network, but also at the epigenome level (Fig. 4e).

### TF sequence binding preferences are similar among family members and conserved through angiosperm evolution

To model TF binding from sequence we applied a “bag-of-k-mers” model^34^ to discriminate TF binding regions from other regions in the genome, which resulted in reliable models for all the TFs (5-fold cross-validation, average accuracy for each TF > 70%) (Fig. 5, Supplementary Fig. 4 and Supplementary Table 10). Using average k-mer weights from the models, we derived a distance matrix among TFs, and summarized TFs relationships (Supplementary Table 11). After removal of singleton families, we observed that for 85% of the TFs families, most of their members (>= 50%) cluster into the same group in a dendrogram (Fig. 5a and Supplementary Fig. 4a).

**Fig. 5.**
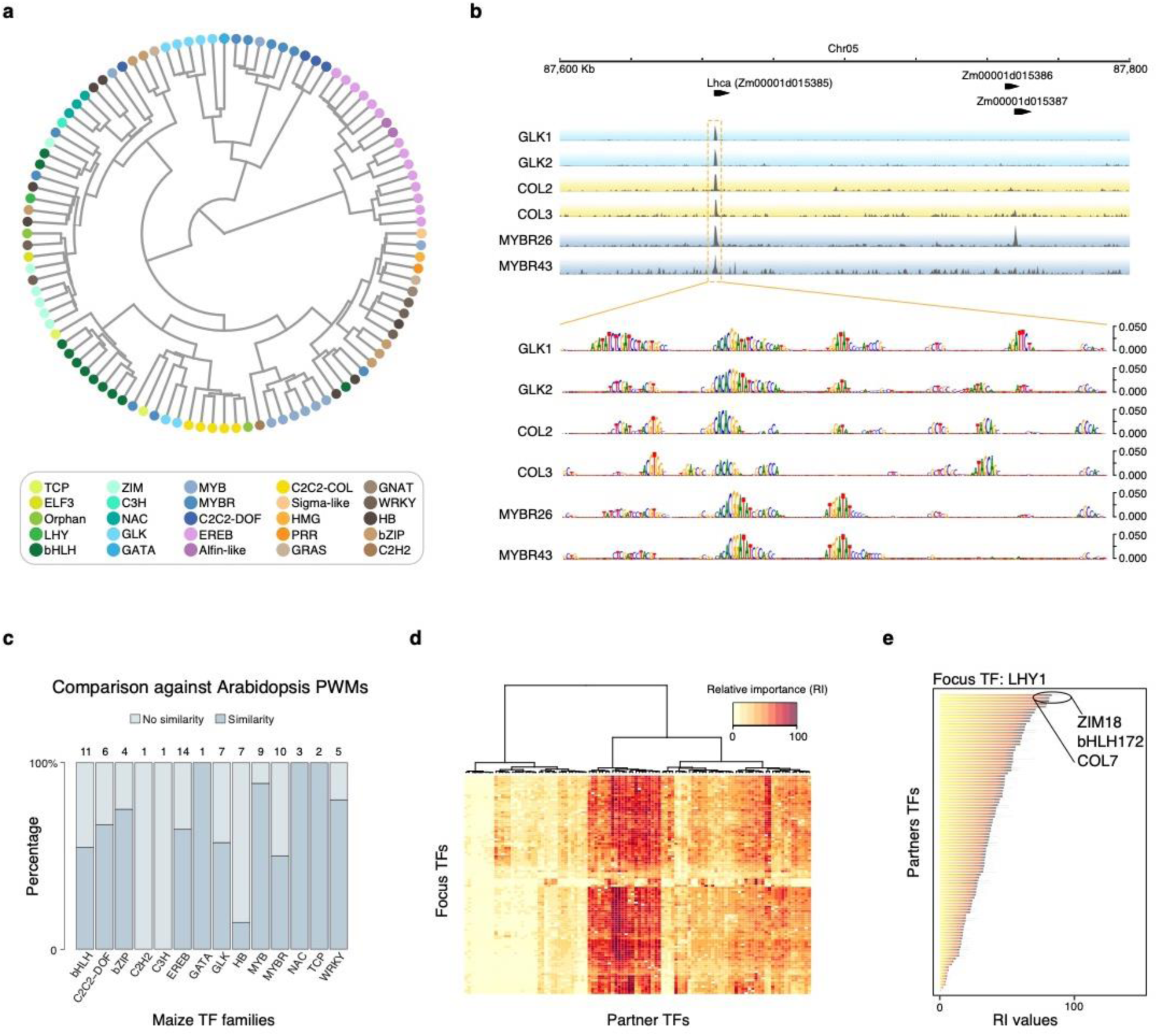
Machine learning models of TF recognition sequence and co-binding. **a**, 104 TF clustered by sequence binding similarity (predicted *k*-mer weights) derived from the sequence models. **b**, Example browser tracks showing the promoter region of *Lhca*, which encodes the light-harvesting complex of photosystem I. Occlusion analysis for TFs targeting *Lhca* promoter showing scores of putative regulatory positions at base pair resolution derived from the sequence models. **c**, Conservation of recognition sequence between arabidopsis and maize TF homologues. Numbers of TFs examined are shown on top, and the TF family names are shown at the bottom. **d**, Cluster of 104 TF based on their relative importance value with co-binding TFs. **e**, Top three co-binding partner TFs of LHY1 based on RI ranking.

This observation prompted us to evaluate whether TF sequence preferences have persisted across angiosperm evolution, as TF protein families are frequently well conserved in plants. Using the top 1% of the predictive *k*-mers for each TF, we examined their similarity to a large collection of Arabidopsis TF binding position weight matrices (PWMs) derived from DAP-seq experiments^35^. After removal of families that did not have a counterpart (or were poorly represented), 50 out of 81 (61%) of the evaluated TFs preferentially match PWMs to their corresponding family in Arabidopsis (P-value < 0.001, Fig. 5c).

Our estimations of conservation through Arabidopsis are likely understated, as some TF families can recognize more than one type of motif, which might not be properly represented in the current databases. For instance, DAP-seq data is not available for the Arabidopsis homologous to *ZmGLK1*, which together with *ZmGLK2* forms an independent cluster from the others GLKs. Yet, our *k*-mer model for *ZmGLK1* (Supplementary Table 11) agrees with the core of a motif (*i*.*e*., “RGAT”, R = A/G) enriched in the promoters of genes up-regulated by Arabidopsis GLK1^30,36^. Overall, the sequence preference conservation suggests a strong constraint over more than 150 million years, which also agrees with the reduced SNP variation at the TF peaks (Fig. 2a).

### TF binding co-localization is context specific

With a genome larger than two billion bp, any given TF (recognition sites avg. 6.8 bp) could have affinity for roughly a third of a million locations across the genome^34,35^. Under this scenario, achieving the observed TF binding specificity would likely require extra cues. We predict that co-binding and combinatorial recognition of *cis*-elements is key for specificity. To test this, we created machine-learning models based on co-localization information to learn non-linear dependencies among TFs using the ENCODE pipeline^5^. To fit a model for each TF (*i*.*e*., focus TF or context), we built a co-localization matrix, by overlapping peaks for the focus-TF with peaks of all remaining TFs (*i*.*e*., partner TF). The co-localization model was aimed to discriminate between the true co-localization matrix and a randomized version of the same^37^. The output of each model is a set of combinatorial rules that can predict TF binding. For each TF, the average of 10 models with independent randomized matrices have area under the receiver operating curve > 0.9 (Supplementary Table 12). This high performance supports the hypothesis that TF co-localization has vast information content to determine binding specificity.

Using the rules derived from the co-localization models, we scored the relative importance (RI) of each partner TF for the joint distribution of the set of peaks for a given context (Fig. 5d and Supplementary Table 13). To obtain a global view, we calculated the average RI of a TF across all focus TFs. We observed that the whole set shows a trend towards medium to low average RI values (*i*.*e*., ≤ 60 RI, more context specific), with fewer TFs predictive for a large number of focus TFs (*i*.*e*., > 60 RI, high-combinatorial potential). For example, among the 104 TFs we examined, LATE ELONGATED HYPOCOTYL (LHY, Zm00001d024546) is the most highly expressed one in the leaf differentiating section^10^. LHY encodes a MYB TF that is a central oscillator in the plant circadian clock^38^, and the its top three predicted partner TFs are ZIM18, bHLH172 and COL7 (Fig. 5e). Although their functions have yet to be characterized in maize, their Arabidopsis homologs are involved in jasmonic acid signaling, iron homeostasis and flowering time regulation, all of which are tightly coupled to the circadian clock^39^.

Clustering according to the RI values revealed a group of TFs that have large RI across many focus TFs (Supplementary Table 13). The large RI score means that a TF is highly predictive of the binding in a specific context. Interestingly, some of them are homologues to Arabidopsis TFs with known roles in hormone signaling pathways (*e*.*g*. ZIM TFs in JA signaling). Some belong to families that are known for protein-protein interactions, such as the MYB-bHLHs and the ZIM-MYB-bHLHs TFs^40–42^, suggesting that these high RI factors could function as to integrate multiple inputs to coordinate transcriptional output. In summary, co-binding could be the key to explain how plant TFs with similar sequence preferences could target different genes and control different biological functions. The co-localization model revealed a large combinatorial space for TF binding sites that likely favors the occurrence of specific combinations, which could facilitate rapid diversification of the regulatory network during speciation.

### Conservation of the TF regulatory interactions among grasses

Finally, we used the machine learning models to investigate how the transcription regulatory network evolved in grasses. To do so we performed ATAC-seq in sorghum and rice, and obtained open chromatin sequences of their maize synteny genes. Next, we inferred network edge conservation based on whether the sequence model of maize TF could predict binding in the open chromatin of the synteny target gene in sorghum and rice (Fig. 6a). For example, we could predict TF binding for 68% of the syntenic open chromatin regions in sorghum. Looking at the network edges from synteny TF to synteny genes, we found that ∼28% of the observed edges in the maize network were conserved to sorghum, and ∼19% were conserved to rice (Fig. 6b).

**Fig. 6.**
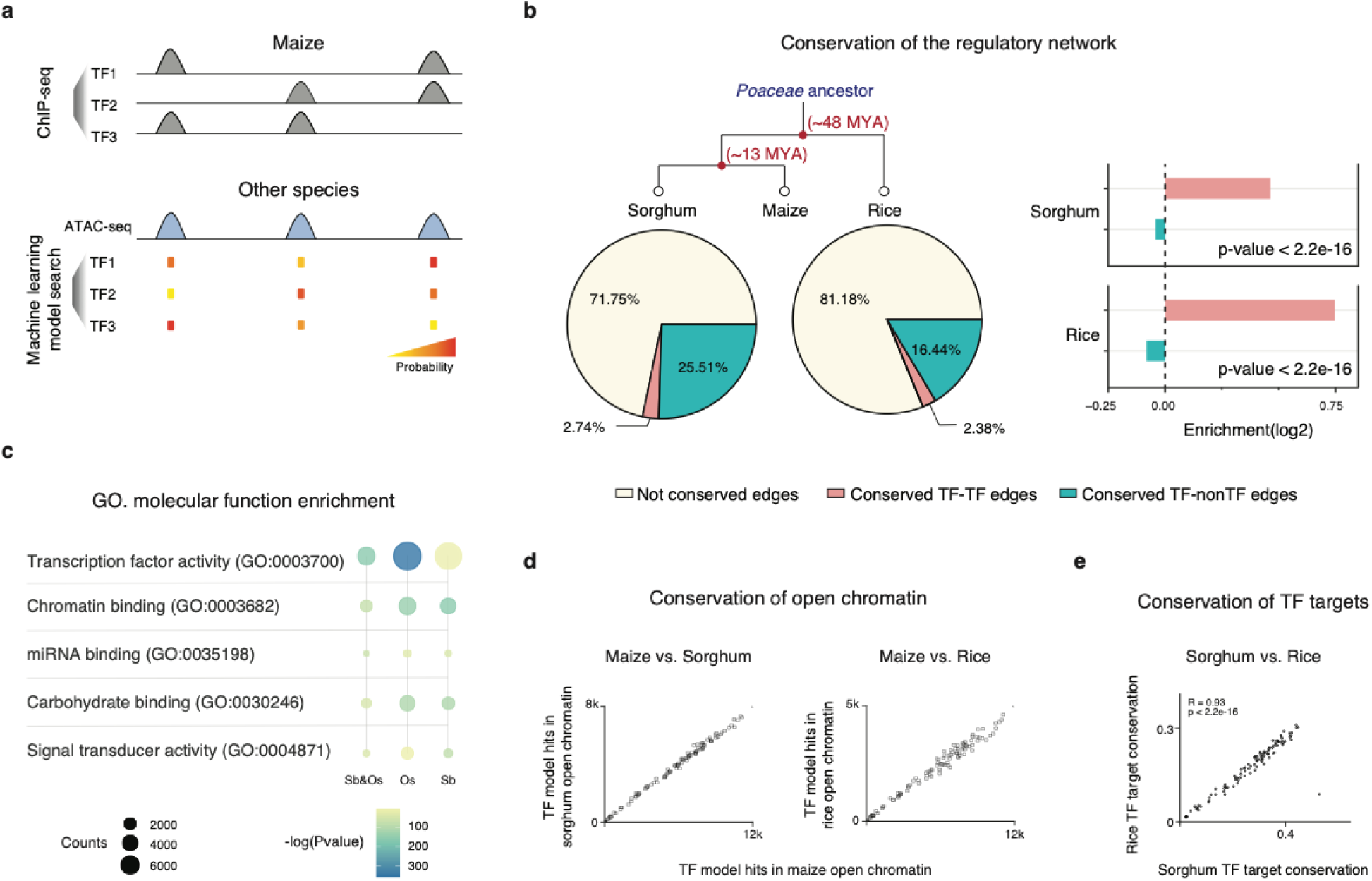
Conservation of the TF regulatory interactions. **a**, Maize TF binding models are used to search the open chromatin regions of maize synteny gene in rice and sorghum, to infer conservation of the regulatory interaction. **b**, Pie chart showing percentage of conerserved edges in sorghum (left) and rice (right). TF to TF edges are more conserved than TF to non-TF gene edges. **c**, GO-term enrichment analysis of the network edges conserved in sorghum, rice and both. **d**, Number of recognition sequences of each TF predicted in maize, sorghum and rice open chromatin. **e**, The axis indicates the percentage of targets conservation of each maize TF in rice and sorghum, inferred from the TF model matches in their synteny target gene open chromatin.

Comparative studies of animal regulatory networks has shown that the core network, which consists of TF-to-TF connections, are frequently more conserved than the rest^43^. If the same is true in plant, we would expect TF genes to be enriched in the set of conserved targets. Hence, we conducted GO enrichment analysis and the result agrees with our expectation, with target nodes enriched for transcription regulatory roles (Fig. 6c). In addition, we also examined the ratio of conserved edges, and found that the TF-to-TF edges were over-represented than TF-to-nonTF edges in both sorghum and rice (Fig. 6b).

Data from modENCODE revealed a strong correlation of the human and mouse homolog TF recognition sites, and the prevalence of TF binding sites in open chromatin is under selection^43^. To test this in plant, we calculated the number of each TF model matches in open chromatin regions in maize, sorghum and rice, and found that they are indeed correlated (Fig. 6d). In addition, for each maize TF, the numbers of conserved targets found in rice and sorghum are also correlated (Fig. 6e), suggesting similar selection pressure during evolution. Taken together, our findings suggest that plants and animals might have adopted a similar strategy to evolve transcription regulatory network, and the rewiring of the network occurs in a hierarchical fashion, with the core network nodes at the top of the transmission of information being more conserved than those at the bottom.

## Discussion

In this study, we attempted to systematically identify the emergent properties of the transcription regulatory network in a plant tissue. Using high-throughput ChIP-seq, we have successfully reconstructed a regulatory network that covers ∼77% of the leaf expressed genes (Fig. 3a). One advantage of its scale-free topology is the increased tolerance to random failures^25^, while the high connectivity hubs are the vulnerabilities. Using chlorophyll biosynthesis and C_4_ photosynthesis as examples (Fig. 4), we showed that mutation of highly connected TFs could indeed disrupt plant growth and development. It also demonstrated how network connectivity data could be used to predict biological functions and identify potential regulators. The observed structural and functional split of the network also indicates how gene expression inside the cell is fine tuned. For example, the presence of multiple TFs across modules to regulate genes in the same pathway suggests intricate modes of actions to coordinate transcription, in contrast to the classic view of a singular or few master regulators.

Our study indicates that organization of plant and animal regulatory networks display a striking level of similarity, despite more than one billion years of evolution and lack of detectable sequence conservation, suggesting this could be of important evolutionary advantage^2,5^. For example, TF co-binding is considered as the most important mechanism of gene regulation in multicellular eukaryotes, and this could be benificial for rapid diversification of their regulatory code. Because from an information standpoint, it enables a large variety of transcriptional outputs using synergistic and combinatorial inputs of a small number of TFs. For instance, taking sets of five distinct TFs (*i*.*e*., the mode of distinct TF binding sites in maize open chromatin regions) from the 104 TF repertoire would generate a ∼91 million possible combinations. In mammalian cells, the high-combinatorial factors, such as MAX and P300, have key roles in enhancer function^44,45^. In our dataset, we also found high-combinatorial TFs (Supplementary Table 13), most of which has not being characterized, and deserve further attention.

The complexity and redundancy of the plant transcription regulation is in agreement with previous studies in other eukaryotic models^*3-6*^, suggesting that these features could provide evolutionary advantages, such as increased tolerance to perturbations, and serving as a reservoir to stored neutral mutations for future usage. Plant researches have been limited to studying either the function of a single gene or perform genome-wide association analysis for everything. The former often resulted in over-simplification, while the latter failed to determine causality. Hence, a bottom-up study towards a systems biology approach is most suited to understand complex and redundant system, and we need to study the system as a whole, not just the individual components.

The large majority of the genome-wide association study hits fall into non-coding regions, with non-coding variants explaining as much phenotypic variation as genic ones. Yet, functional interpretation of non-coding variation is a challenge. The extensive TF bidning data we provided should greatly facilitate future analysis into the organization and regulation of plant genome, and the presented TF binding and co-binding models could be integrated into pipelines to predict effects of non-coding variants, both common and rare, on TF binding, to pinpoint causal sites. As the possibility of being able to predict and generate novel variations not seen in nature could fundamentally change future plant breeding.

## Data availability

Sequencing data have been deposited in the NCBI SRA database under the accession number PRJNA518749. Processed data have been deposited in the NCBI GEO database under the accession number GSE137972, and will be incorporated into the future MaizeGDB release.

## Code availability

Code to reproduce analyses is available at https://bitbucket.com/p_transcriptionfactors/analysis.

## Acknowledgements

We thank B. Du, X. Wang, R. Foo, J. Li and M. Huang for technical support, G. Ramstein for advice in the analysis of GWAS data, and D. Grierson and S. Miller for proofreading the manuscript.

## Author contributions

S. Zhong, E.S. Buckler, and P. Li designed and supervised the research; X. Tu. developed techniques, performed experiments and processed the raw data with assistance from D. Tzeng, P. Chu and X. Dai. M.K. Mejia-Guerra designed and performed computational analysis, J.A.Valdes Franco performed population genetics analysis; M.K. Mejia-Guerra and S. Zhong wrote the paper. All the authors read the paper and agree with the final version.

## Materials & Correspondence

Biological material requests should be addressed to S.Z. and P. L., and requests for codes and analysis should be addressed to M.K.M.

## Supplementary Information

### Supplementary Figures S1-S7

### Supplementary Tables S1-S13

